# TumorFusions: an integrative resource for reporting cancer-associated transcript fusions in 33 tumor types

**DOI:** 10.1101/162180

**Authors:** Xin Hu, Qianghu Wang, Floris Barthel, Ming Tang, Samirkumar Amin, Kosuke Yoshihara, Frederick M. Lang, Soo Hyun Lee, Siyuan Zheng, Roel G.W. Verhaak

## Abstract

Fusion genes, particularly those involving kinases, have been demonstrated as drivers and are frequent therapeutic targets in cancer^1^. Here, we describe our results on detecting transcript fusions across 33 cancer types from The Cancer Genome Atlas (TCGA), totaling 9,966 cancer samples and 648 normal samples^2^. Preprocessing, including read alignment to both genome and transcriptome, and fusion detection were carried out using a uniform pipeline^3^. To validate the resultant fusions, we also called somatic structural variations for 561 cancers from whole genome sequencing data. A summary of the data used in this study is provided in **Table S1**. Our results can be accessed per our portal at http://www.tumorfusions.org.

---

We identified 56,198 and 3,838 candidate fusions in 9,966 cancer and 648 non-neoplastic samples, respectively. After applying stringent filters controlling for sequence similarity of the partner genes, transcriptional allelic fraction, dubious junctions, germline events, and presence in non-neoplastic tissue, we obtained 20,731 high confident fusion events (**Table S1**), 54% of which were supported by at least one DNA breakpoint near the fusion junction per the matching Affymetrix SNP6 DNA copy number data profile. Frequent germline fusions between adjacent genes such as between *CRHR1* and *KANSL1* on chr17q (n=36, 6%) or *TFG* and *GPR128* (n = 9, 1%) on chr3q12.2 were frequently associated with focal copy number changes and were likely the result of germline polymorphisms^4^. We compared fusions with somatic structural variations (SVs) detected using whole genome sequencing from 561 cancer samples, which associated 1,679 of 2,585 fusions (65%) to SVs. Of these, the majority were translocations (50%), followed by transcript fusions as a result of deletions (20%), inversions (20% or duplication (10%) (**Figure S1**). While 57% of fusions (n=962) mapped to a single SV, 21% (n=348) were associated with three or more SVs. Such fusion events likely resulted from complex DNA rearrangement events. We found a significant enrichment for chromosome arm 12q fusions in sarcoma (adjusted p<0.001, Chi-square test) (**Figure S2**), seen previously in glioblastoma^5^. The 12q13-15 growth factor signaling gene *FRS2* was frequently involved as the 5’ partner (**Figure S3**), nominating *FRS2* as a relevant target in this disease^6^. Pertaining to the number of sarcomas with a complex 12q (21%), this cancer showed enrichment for cases with an excessive number of transcript fusions (9% in sarcoma vs 2% in other cancers, p= 1.78e-21, Chi-square test; **Figure S4**).

The majority of the 20,731 fusions were singletons (n = 17,238, 83.2%). Amongst the 1,205 recurrent fusions, 850 were found in only two cases. These data suggest that gene fusions are frequently not selected for but represent collateral DNA rearrangement damage. The most frequent recurrent fusions were lineage specific, such as *TMPRSS2-ERG* in prostate cancer, *CCDC6-RET* in thyroid cancer, and *PML-RARA* and *CBFB-MYH1* in acute leukemia. In contrast *FGFR3-TACC3* (n = 36), *PTPRK-RSPO3* (n = 9) and *EML4-ALK* (n = 7) were found across multiple originating tissues (**Figure 1A**). Breaking fusions down into their separate gene partners, we found that other than *TMPRSS2* and *ERG*, recurring fusion partner genes were usually found in more than one tissue of origin (**Figure 1B**). *TRK* fusions, targeting of which by larotrectinib has recently shown promising clinical efficacy^7^, were found in 28 cases across 11 cancer types. Other fusions with potential clinical relevance included *BRAF* associated fusions in 30 cases from 11 cancer types, for which sorafenib may provide a therapeutic advantage^8^; *MET* fusions in 20 cases from ten tumor types which may respond to crizotinib^9^, and *ROS1* fusions in six cases of lung adenocarcinoma/glioblastoma^10^. In aggregate 7,470 genes were found in fusions in more than one sample, as either 5’ or 3’ partner gene, representing about 30% of the annotated genes in the human genome.

**Figure 1A.**
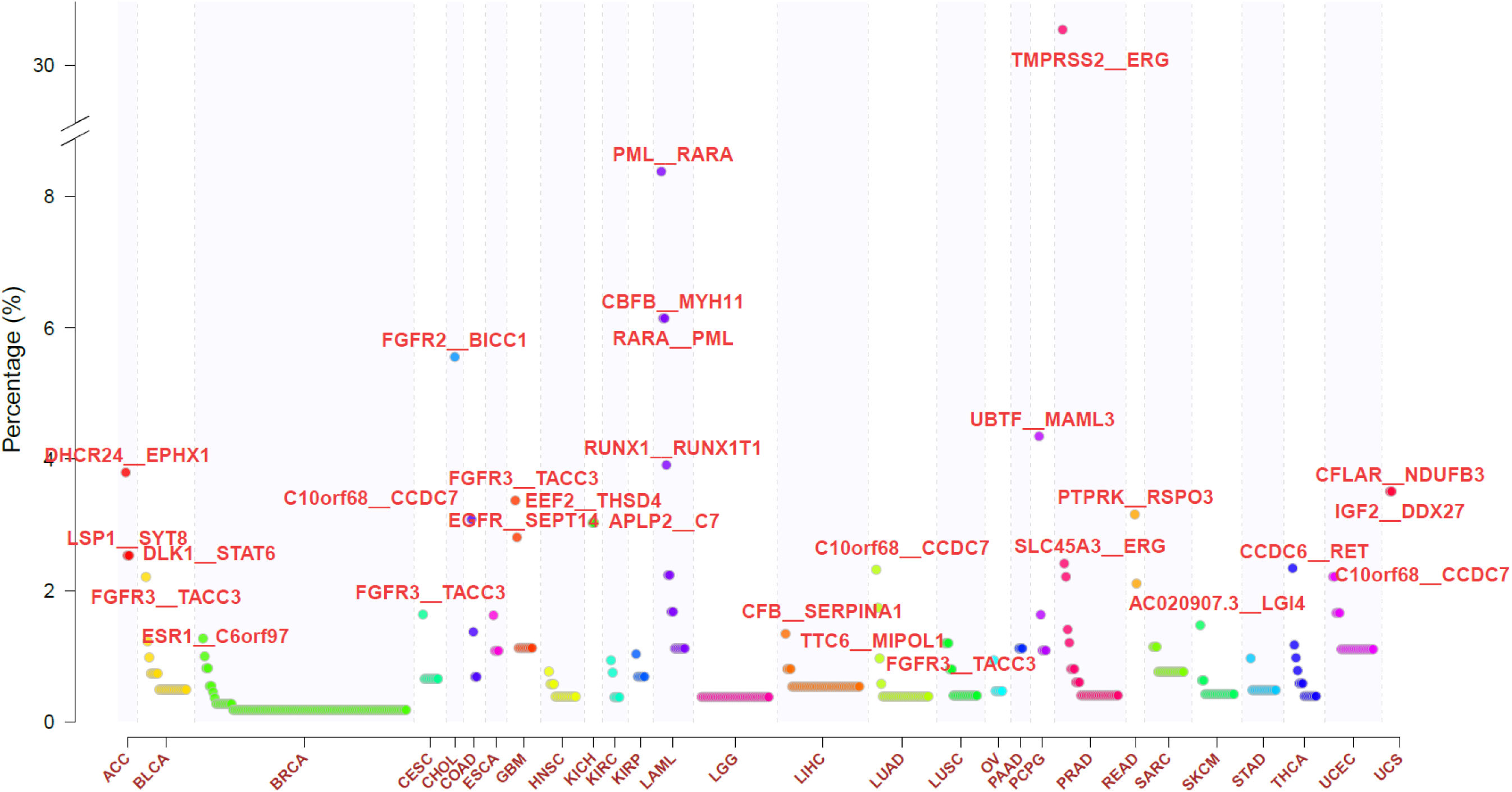
Frequency of recurrent fusions across 33 cancer types. Y axis represents percentage of cohort wherein the fusion is found. Only recurrent fusions are shown in the figure.

**Figure 1B.**
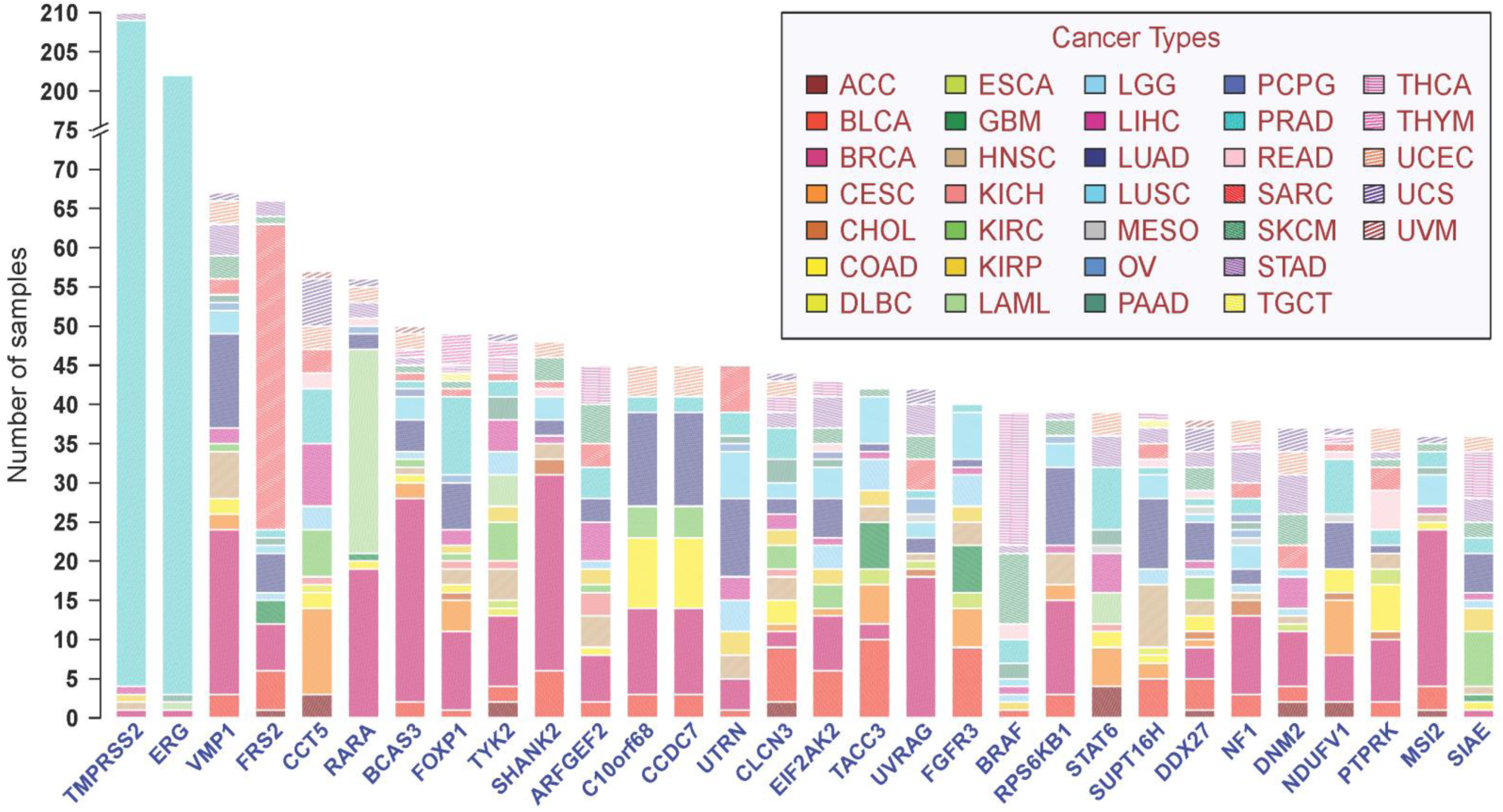
Top frequent partner genes in recurrent fusion transcripts across 33 cancer types. Y-axis represents frequency of partner genes in the pan-cancer cohort.

To prioritize potential functional events we intersected the fusion list with tumor suppressors, oncogenes, kinases, and epigenetic modifiers. We identified 763 fusions involving a kinase, of which 341 retained their kinase domains. The most frequent, novel kinase fusion was *TMEM87B-MERTK* found in 7 cases. Mutations in *MERTK* has been associated with retinal degeneration as well as various human cancers^11^. We evaluated the functional relevance of fusion candidates by computing a neighboring gene network based centrality metric^12^. Known driver fusions had significantly higher centrality scores than other fusions (p<2.2e-16, t-test; **Figure S5**). Notable novel recurrent fusions with high centrality scores include *ERC1*-*RET* (n=3), *ERBB2*-*PPP1R1B* (n=4), and *KLK2-FGFR2* (n=3).

We have built a fusion portal to facilitate broad access to this resource (www.tumorfusions.org). Users can search the portal by gene, fusion, sample, or browse it by cancer type, and annotation is provided for each fusion candidate using the comprehensive and rich data portfolio from TCGA. The portal links each partner gene of a fusion to its copy number and mutational pattern, a functionality that allows users to visually assess the association of the fusion with copy number, and the functional significance of the partner genes in the cancer type. The portal also annotates each partner gene as to whether they are cancer genes (oncogene/tumor suppressor), epigenetic modifiers, kinases, or whether they lose post translational modification sites (phosphorylation, ubiquitylation). For each fusion, the portal provides images depicting the exon expression in relation to the junction, relevant gene annotation and the expression of the two partner genes (**Figure S6)**, both of which are useful for evaluating the functional impact of the fusion. Fusion transcript represent a class of infrequent yet often targetable somatic alteration. Technological advances allow testing the oncogenicity of these events in a high throughput manner^13^ and clinical trial design is evolving towards adaptive design across cancer baskets^14^, supported by our ongoing characterization of transcript fusions.

## ACKNOWLEDGEMENTS

This study is in whole or partially based upon data generated by The Cancer Genome Atlas project established by the NCI and NHGRI. Information about TCGA and the investigators and institutions that constitute the TCGA research network can be found at http://cancergenome.nih.gov. The authors thank Jinzhen Chen and Boyd Sally for their IT support. This project is supported by grants from the National Institutes of Health R01CA190121 to R.G.W.V., P50CA127001 to Z.S. and R.G.W.V. and the Cancer Prevention and Research Institute of Texas (CPRIT) (R140606) to R.G.W.V. This work was supported by Cancer Center Support Grants P30CA16672 and P30CA034196.

## Supplementary Methods

### Data resources

TCGA DNA and RNA sequencing data were downloaded from Cancer Genomics Hub (CGHub, https://cghub.ucsc.edu). Copy number segmentation data and gene expression data were downloaded from Firehose (https://gdac.broadinstitute.org/). Somatic mutation data were downloaded from UCSC Xena repository (https://xenabrowser.net/datapages/?cohort=TCGA%20Pan-Cancer%20(PANCAN)). Genomic Variants database was retrieved from http://dgv.tcag.ca/dgv/app/home. All data used in this study were summarized in **Table S1**. We excluded 41 samples from the 689 normal samples because they clustered with tumor samples in unsupervised hierarchical clustering. The clustering was done within each cancer type using expression of all genes and ward’s method. The resulting panel of normal samples (n=648) were subjected to the same fusion detection pipeline and were used as controls to filter out potential germline fusion events and artifacts.

### Identification of fusion transcripts

We applied PRADA ^3^ to all RNAseq samples for data preprocessing and fusion calling. In brief, RNA sequencing reads were aligned to a composite reference consisting of both genome (hg19) and transcriptome (Ensembl 64), followed by a remapping step that aligns transcriptome coordinates to the reference genome^15^. GATK best practices were implemented in the pipeline, including marking duplication and base quality recalibration. More information about PRADA can be found at http://bioinformatics.mdanderson.org/main/PRADA:Overview.

PRADA detects fusion transcripts based on discordant read pairs (reads mapping to different protein-coding genes) and junction spanning reads (reads mapping to the exon–exon junctions). We required at least two discordant read pairs and one junction spanning read to call a fusion candidate. All fusion candidates were collected and were subject to additional filtering. The filters were described as follows: (1) candidates observed in normal controls were removed; (2) candidates with highly similar partners in sequence (blastn e-value≤0.001) were removed; (3) candidates with low transcriptional allelic fraction were removed (TAF, minimum 0.01 for both partner genes); (4) candidates with very promiscuous partner genes were removed (the Partner Gene Variety filter, see below); (5) Candidates with identical junctions in more than 15 samples were removed; (6) candidates with supporting reads mapped disproportionately to sense and antisense strands were removed. Transcriptional allelic fraction (TAF) was calculated as the ratio of fusion supporting junction spanning reads to the total number of reads spanning the junction involved in the fusion. Partner Gene Variety (PGV) was defined as the number of unique chromosomal arms where the partner genes were found. A higher PGV suggests a gene was found to fuse with more partner genes in a cancer lineage. For genes with PGV greater than 10, we used permutations (n=100,000) to model the background distribution of the random chances of obtaining the observed PGV (empirical p value). We removed fusions with empirical p value less than 0.001%. For filter (6), we hypothesized that ratio of sense and antisense strand mapping reads was proportional to the distance from the start of the fusing transcript to the junctions of the two partner genes. Since lower coverage and short distance may confound this ratio, we limited our filtering to fusions with more than 100 spanning reads and such distance more than 500 base pairs. We removed fusions that had this ratio greater than 100.

To establish a positive control fusion list, we integrated three resources including Mitelman^16^, ChimerPub^17^, and Cosmic fusions^18^. Fusions reported in all three independent references were curated as a list of known fusions (n=321). Of these 321, 38 fusions were detected in our data set reflecting 359 instances in total.

### Validation of fusion transcripts through integrating structure variants and copy number changes

For cases where both copy number profile and gene fusion were available, we aligned fusion junctions with copy number breakpoints. We allowed a 100 Kb window to the expected orientation for both partner genes when searching array based copy number data.

We detected structural variants (SVs)^19^ from whole genome sequencing (WGS) data using Speedseq with default parameters. We filtered SVs requiring more than 3 supporting reads, i.e. at least one split read and one discordant read pair. For fold-back inversions (BND on the same chromosome) we required more than 9 supporting reads. We removed SVs with breakpoints falling in low-complexity regions (e.g. repeat region DNA), or stacking across different tumor types. We further removed SVs where the flanking 100 bp of the two breakpoints share high sequence similarity (blastn E-value > 0.0001). Germline events were filtered out by comparing with matched normal samples.

We scanned the intersection between the edge of confident interval from the supported structure variants including large fragment duplication, deletion, insertion and inversion and truncated intron region flanking the junction upon fusion events. We assigned two partner genes into three groups based on their relevant location of break points to adjacent break point of structure variants. High confidence group was defined when a break point of structure variants fell into the immediate intron of the fusing exon for both partner genes; low confidence group was defined when a break point of structure variants fell between the fusion junction and the start or end of the partner gene depending on the fusion orientation, or fell into the 100K window from the corresponding gene boundary; Intermediate confidence group was defined when one partner gene met criteria of high confidence group and the other met that of the low confidence group. For those fusion pairs with only one junction points supported by structure variants, we assigned as one-sided.

### Exons and transcription expression analysis of fusions

Exons and transcripts expression of fusion partners are retrieved from normalized RSEM value of level3 RNA-seq from Firehose (https://gdac.broadinstitute.org/). We performed Z score transformed expression level across all samples in each cancer type to plot exon expression heatmaps.

### Fusion centrality analysis

Fusion transcript centrality score was calculated based on domain-based fusion model using default parameters (https://bmsr.usc.edu/software/targetgene/), to predict the oncogenic driver in which partner genes act as hubs in a cancer pathway network^12^. Fusions with centrality score > 0.37 were considered as potential drivers.

## REFERENCES

1. Mertens, F., Johansson, B., Fioretos, T. & Mitelman, F. The emerging complexity of gene fusions in cancer. Nat Rev Cancer 15, 371–381 (2015).

2. Yoshihara, K. et al. The landscape and therapeutic relevance of cancer-associated transcript fusions. Oncogene 34, 4845–4854 (2015).

3. Torres-Garcia, W. et al. PRADA: pipeline for RNA sequencing data analysis. Bioinformatics 30, 2224–2226 (2014).

4. Iafrate, A. J. et al. Detection of large-scale variation in the human genome. Nat Genet 36, 949–951 (2004).

5. Zheng, S. et al. A survey of intragenic breakpoints in glioblastoma identifies a distinct subset associated with poor survival. Genes Dev 27, 1462–1472 (2013).

6. Wu, Y., Chen, Z. & Ullrich, A. EGFR and FGFR signaling through FRS2 is subject to negative feedback control by ERK1/2. Biol Chem 384, 1215–1226 (2003).

7. Garber, K. In a major shift, cancer drugs go ‘tissue-agnostic’. Science 356, 1111–1112 (2017).

8. Ross, J. S. et al. The distribution of BRAF gene fusions in solid tumors and response to targeted therapy. Int J Cancer 138, 881–890 (2016).

9. International Cancer Genome Consortium PedBrain Tumor, P. Recurrent MET fusion genes represent a drug target in pediatric glioblastoma. Nat Med 22, 1314–1320 (2016).

10. Drilon, A. et al. Safety and Antitumor Activity of the Multitargeted Pan-TRK, ROS1, and ALK Inhibitor Entrectinib: Combined Results from Two Phase I Trials (ALKA-372-001 and STARTRK-1). Cancer Discov 7, 400–409 (2017).

11. Parinot, C. & Nandrot, E.F. A Comprehensive Review of Mutations in the MERTK Proto-Oncogene. Adv Exp Med Biol 854, 259–265 (2016).

12. Wu, C.C., Kannan, K., Lin, S., Yen, L. & Milosavljevic, A. Identification of cancer fusion drivers using network fusion centrality. Bioinformatics 29, 1174–1181 (2013).

13. Lu, H. et al. Engineering and Functional Characterization of Fusion Genes Identifies Novel Oncogenic Drivers of Cancer. Cancer Res 77, 3502–3512 (2017).

14. Redig, A.J. & Janne, P.A. Basket trials and the evolution of clinical trial design in an era of genomic medicine. J Clin Oncol 33, 975–977 (2015).

15. Berger, M.F. et al. Integrative analysis of the melanoma transcriptome. Genome Res 20, 413–427 (2010).

16. Mitelman, F., Johansson, B. & Mertens, F. The impact of translocations and gene fusions on cancer causation. Nat Rev Cancer 7, 233–245 (2007).

17. Lee, M. et al. ChimerDB 3.0: an enhanced database for fusion genes from cancer transcriptome and literature data mining. Nucleic Acids Res 45, D784–D789 (2017).

18. Forbes, S.A. et al. COSMIC: exploring the world’s knowledge of somatic mutations in human cancer. Nucleic Acids Res 43, D805–811 (2015).

19. Chiang, C. et al. SpeedSeq: ultra-fast personal genome analysis and interpretation. Nat Methods 12, 966–968 (2015).

